# Metaxin 3 is a Highly Conserved Vertebrate Protein Homologous to Mitochondrial Import Proteins and GSTs

**DOI:** 10.1101/813451

**Authors:** Kenneth W. Adolph

## Abstract

Metaxin 3 genes are shown to be widely conserved in vertebrates, including mammals, birds, fish, amphibians, and reptiles. Metaxin 3 genes, however, are not found in invertebrates, plants, and bacteria. The predicted metaxin 3 proteins were identified by their homology to the metaxin 3 proteins encoded by zebrafish and *Xenopus* cDNAs. Further evidence that they are metaxin proteins was provided by the presence of GST_N_Metaxin, GST_C_Metaxin, and Tom37 protein domains, and the absence of other major domains. Alignment of human metaxin 3 and human metaxin 1 predicted amino acid sequences showed 45% identities, while human metaxin 2 had 23% identities. These results indicate that metaxin 3 is a distinct metaxin. A wide variety of vertebrate species—including human, zebrafish, *Xenopus*, dog, shark, elephant, panda, and platypus—had the same genes adjacent to the metaxin 3 gene. In particular, the thrombospondin 4 gene (*THBS4*) is next to the metaxin 3 gene (*MTX3*). By comparison, the thrombospondin 3 gene (*THBS3*) is next to the metaxin 1 gene (*MTX1*). Phylogenetic analysis showed that metaxin 3, metaxin 1, and metaxin 2 protein sequences formed separate clusters, but with all three metaxins being derived from a common ancestor. Alpha-helices dominate the predicted secondary structures of metaxin 3 proteins. Little beta-strand is present. The pattern of 9 helical segments is also found for metaxins 1 and 2.

## INTRODUCTION

The first metaxin 3 to be identified and investigated was zebrafish metaxin 3 (Adolph, 2004). cDNAs encoding zebrafish metaxin 3, along with cDNAs for metaxins 1 and 2, were sequenced and properties of the encoded proteins were deduced. Zebrafish (*Danio rerio*) was studied because it is an important model system for research into vertebrate development, molecular biology, and genomics (Choudhuri et al., 2017; Cagan et al., 2019; Simone et al., 2018). The zebrafish metaxin 3 cDNA was found to encode a protein of 313 amino acids (MW 35,208), which has 40% amino acid identities compared to zebrafish metaxin 1 and 26% identities compared to metaxin 2. In addition to amino acid sequence homology, the predicted zebrafish metaxin 3 protein sequence was defined by the presence of GST_N_Metaxin and GST_C_ Metaxin domains. A common ancestry for the three metaxins was revealed by phylogenetic tree analysis. Alpha-helical segments were found to be the predominant feature of zebrafish metaxin 3 predicted secondary structure.

Metaxin 3 cDNAs were also identified and sequenced for *Xenopus laevis* (Adolph, 2005). This amphibian is a classic model system for studies of vertebrate early development, the cell cycle, and the cytoskeleton (Blow and Laskey, 2016; Bier and De Robertis, 2015; Harland and Grainger, 2011). In comparing the deduced protein sequence of *Xenopus* metaxin 3 with zebrafish metaxin 3, 55% amino acid identities were observed, demonstrating the high degree of conservation of the metaxin 3 protein. That *Xenopus* metaxin 3 is a metaxin 3 protein was supported by (1) the presence of characteristic GST_Metaxin domains, (2) phylogenetic trees showing that *Xenopus* metaxin 3 and other metaxin 3 proteins form a separate group, and by (3) a pattern of alpha-helical secondary structure conserved among metaxin 3 proteins.

The metaxin 1 gene was first identified as a mouse gene (*Mtx1*) located on chromosome 3 between thrombospondin 3 (*Thbs3*) and glucocerebrosidase (*Gba*) genes (Bornstein et al., 1995). The metaxin 1 protein of 317 amino acids was found to be essential for embryonic development, and is ubiquitously expressed in mouse tissues. Experimental evidence revealed that metaxin 1 is a protein of the outer mitochondrial membrane involved in protein import into mitochondria (Armstrong et al., 1997).

Human tissues express a homologous protein (MTX1). The gene (*MTX1*) is located at cytogenetic location 1q21 and is flanked by thrombospondin 3 (*THBS3*) and glucocerebrosidase 1 (*GBA1*) genes. The thrombospondins are extracellular glycoproteins that mediate cell-to-matrix and cell-to-cell interactions (Adams and Lawler, 2011; Mosher and Adams, 2012). Mutations in the glucocerebrosidase gene can result in a deficiency of the enzyme and cause Gaucher disease. Gaucher disease is a progressive, inherited condition caused by the build-up of glucocerebroside in many organs and tissues of the body (Mistry et al., 2017). Understanding the genetic basis of the disease is complicated by the close proximity of the *GBA1, MTX1*, and *THBS3* genes, and by the presence of *MTX1* and *GBA1* pseudogenes at the same genetic locus. Recombination and mutation in this region could have a modifying role in Gaucher disease (Tayebi et al., 2003; La Marca et al., 2004).

Mutations in the glucocerebrosidase gene are also associated with an increased genetic risk for the development of Parkinson disease (Do et al., 2019). Glucocerebrosidase and alpha-synuclein, a protein that forms amyloid deposits in brain cells, may interact either directly or indirectly. However, the symptoms of Parkinson disease are not observed in the majority of patients with *GBA1* mutations.

Metaxin 2 was initially identified in the mouse as a protein that interacts with metaxin 1 in the mitochondrial outer membrane (Armstrong et al., 1999). A complex of metaxins 1 and 2 might therefore have a role in protein import into mitochondria. A metaxin 2 cDNA coded for a protein (263 amino acids) with 29% amino acid sequence identities to mouse metaxin 1. A human homolog is on chromosome 2 at 2q31.2. The human gene is not adjacent to thrombospondin or glucocerebrosidase genes, but is next to a cluster of homeobox genes (*HOXD1, D3, D4, D8, D9, D10, D11, D12, D13*) which code for transcription factors.

Protein import into mitochondria has been more extensively studied in yeast and other fungi (Pfanner et al., 2019; Wiedemann and Pfanner, 2017; Neupert, 2015). For example, the insertion of beta-barrel membrane proteins into the mitochondrial outer membrane involves the TOM complex (translocase of the outer mitochondrial membrane complex) and the SAM complex (sorting and assembly machinery complex). A conserved protein domain of one of the component proteins, the TOM37 domain, is also found in the metaxin proteins (see Results and Discussion), providing a further indication of the role of the vertebrate metaxins in transporting proteins across the outer mitochondrial membrane.

Very little is known about metaxin 3 genes and proteins. The only publications that mention metaxin 3 are the cDNA studies of zebrafish and *Xenopus* (Adolph, 2004, 2005). For this reason, metaxin 3 proteins of a wide variety of vertebrates have been investigated. The proteins are the predicted proteins in the NCBI databases primarily derived from genome sequencing. The metaxin 3 gene was found to be present in a wide variety of vertebrate species. Fundamental properties of the encoded proteins were determined, and confirm the high degree of conservation of the metaxin 3 genes and proteins.

## MATERIALS AND METHODS

The conserved protein domains included in Figure 1 (GST_N_Metaxin, GST_C_Metaxin, Tom37) were identified using the CD_Search tool of the NCBI (www.ncbi.nih.gov/Structure/cdd/wrpsb.cgi; Marchler-Bauer et al., 2017). For the amino acid sequence alignments in Figure 2, the GlobalAlign program was used to compare two metaxin sequences along their entire lengths (https://blast.ncbi.nlm.nih.gov/BlastAlign; Needleman and Wunsch, 1970; Altschul et al., 1997). Neighboring genes (Figure 3) were identified from the results of Protein BLAST searches with metaxin 3 sequences and the related information on “Gene neighbors” (https://blast.ncbi.nlm.nih.gov/Blast.cgi). The Genome Data Viewer and, earlier, the Map Viewer genome browsers were also used to identify adjacent genes (www.ncbi.nlm.nih.gov/genome/gdv).

**Figure 1.**
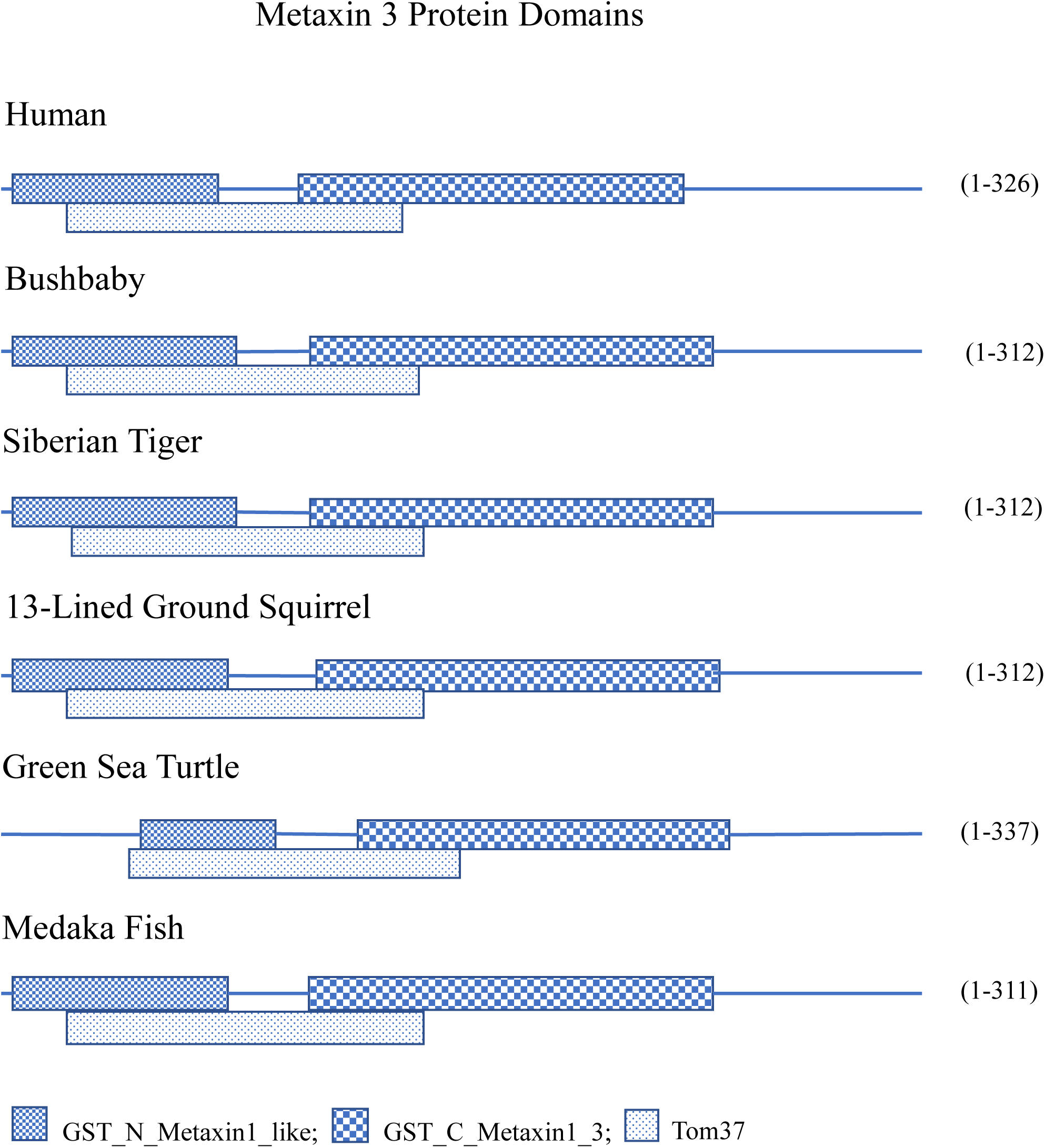
Protein domain structure of metaxin 3. Three major conserved domains are present in each of the examples shown: a GST_N_Metaxin1_like domain, a GST_C_Metaxin1_3 domain, and a Tom7 domain. The examples represent primates (human, bushbaby), a carnivore (Siberian tiger), a rodent (13-lined ground squirrel), a reptile (green sea turtle), and a fish (medaka). The domains are highly conserved in all of the examples shown, and in other metaxin 3 proteins. Metaxin 1 and 2 proteins also have GST_N_Metaxin, GST_C_Metaxin, and Tom37 domains. But the percentages of amino acid identities show that metaxin 3 is a protein distinct from metaxins 1 and 2. Tom37 is a yeast protein of the outer mitochondrial membrane involved in the import of beta-barrel proteins into mitochondria. Its presence is an indication that the metaxins also function in protein import into mitochondria.

**Figure 2.**
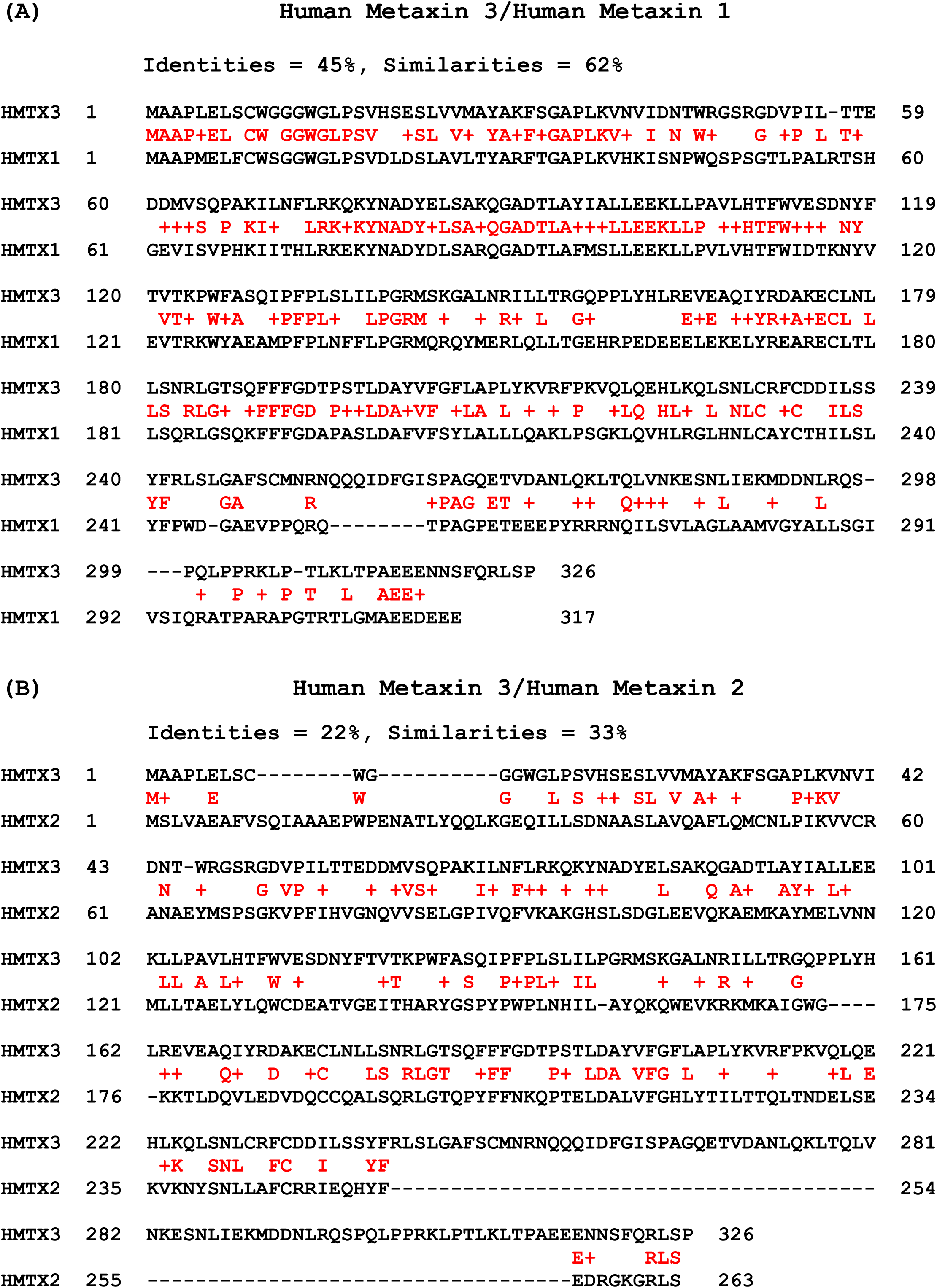
Alignment of the amino acid sequence of human metaxin 3 with human metaxins 1 and 2. (A) The alignment of human metaxin 3 (326 aa) with metaxin 1 (317 aa) shows 45% identical residues and 62% similar residues. The identical and similar amino acids are between the metaxin 3 and metaxin 1 sequences and shown in red. (B) Alignment of human metaxin 3 with human metaxin 2 (263 aa) shows a lower degree of homology, with identities = 22% and similarities = 33%. The identities and similarities are shown in red between the sequences. The percent identities in (A) and (B) demonstrate that metaxin 3 is a protein distinct from metaxins 1 and 2. The results were obtained using the NCBI BLAST Global Align function (Needleman-Wunsch) to compare two sequences along their entire lengths.

**Figure 3.**
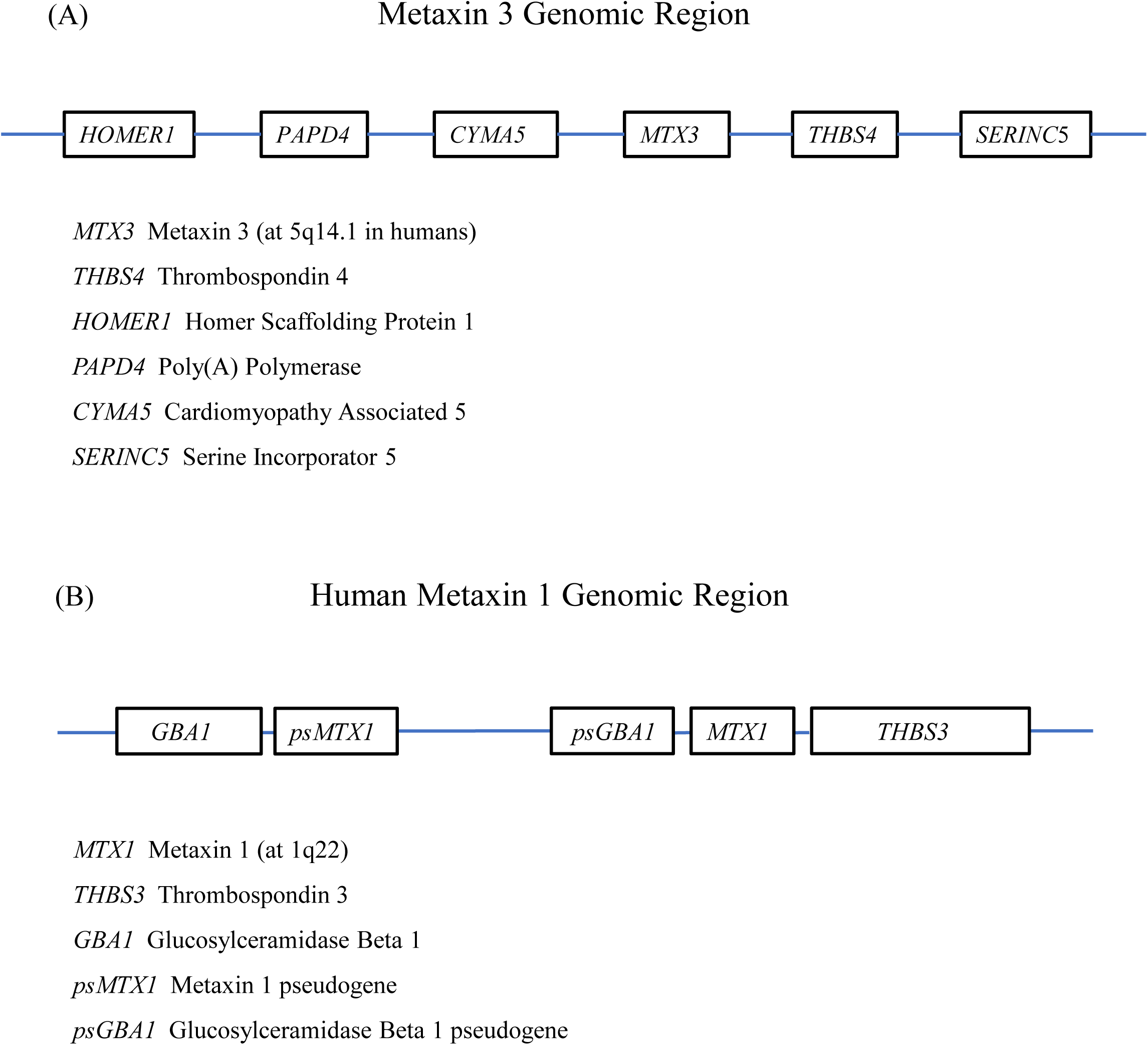
Neighboring genes of metaxin 3 and metaxin 1. (A) includes the genes adjacent to the metaxin 3 gene for a wide variety of vertebrates. The order of the genes is shown, but the distances between the genes are not to scale. For human metaxin 3, the genomic region is on chromosome 5, at 5q14.1. In (B), the immediately adjacent genes for human metaxin 1 at 1q22 are shown. The sizes of the genes and distances between the genes are approximately to scale. Pseudogenes of the *MTX1* gene and *GBA1* gene are a prominent feature of this genomic region. In (A), a thrombospondin gene (*THBS4*) is next to the metaxin 3 gene, and in (B), a thrombospondin gene (*THBS3*) is next to the metaxin 1 gene. But the other neighboring genes are different.

The phylogenetic tree in Figure 4 was generated with the COBALT multiple sequence alignment tool (www.ncbi.nlm.nih.gov/tools/cobalt; Papadopoulos and Agarwala, 2007). The predicted secondary structures of metaxin 3 proteins (Figure 5) were determined with the PSIPRED secondary structure prediction server (bioinf.cs.ucl.ac.uk/psipred/: Jones, 1999). The PHOBIUS program for prediction of transmembrane topology was used to investigate the presence of transmembrane helices (www.ebi.ac.uk/Tools/pfa/phobius/; Madeira et al., 2019). The TMHMM prediction server was also used (www.cbs.dtu.dk/services/TMHMM/; Krogh et al., 2001). Whether metaxin 3 proteins have signal peptides was investigated with the SignalP program (www.cbs.dtu.dk/services/SignalP/; Armenteros et al., 2019)

**Figure 4.**
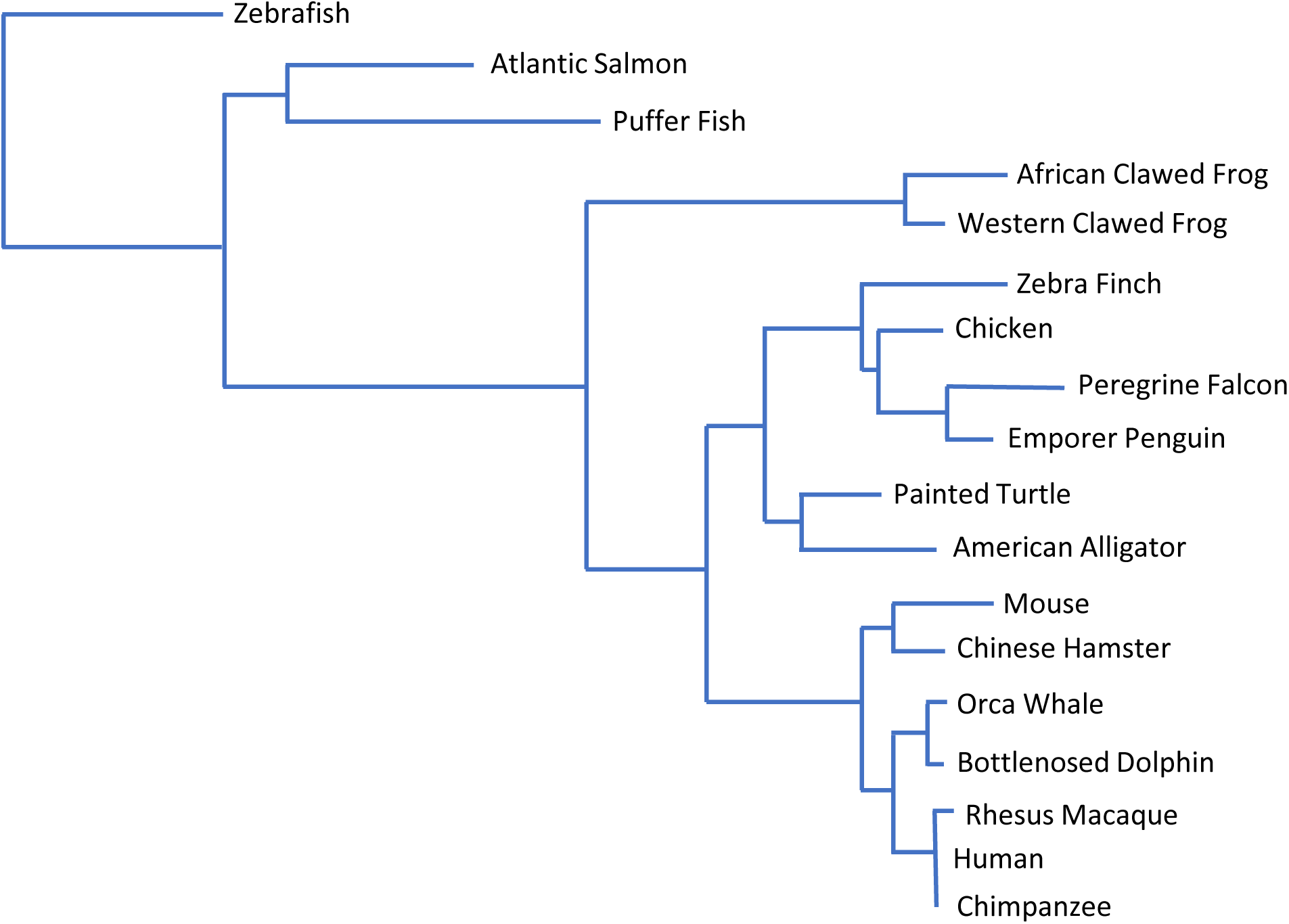
Phylogenetic analysis of metaxin 3 proteins. The phylogenetic tree shows that the metaxin 3 proteins of a selection of diverse vertebrates evolved from a common ancestor. Zebrafish and other fish diverged first, followed by amphibians and then birds, reptiles, and mammals. Similar vertebrates are seen to have similar metaxin 3 sequences. For example, the mammalian metaxin 3 proteins (human, chimpanzee, macaque, dolphin, whale, hamster, mouse) form a distinct cluster, as do fish, amphibians, birds, and reptiles. The analysis was carried out with the COBALT multiple sequence alignment tool (NCBI) using the Neighbor Joining method. The branch lengths are proportional to the evolutionary change.

**Figure 5.**
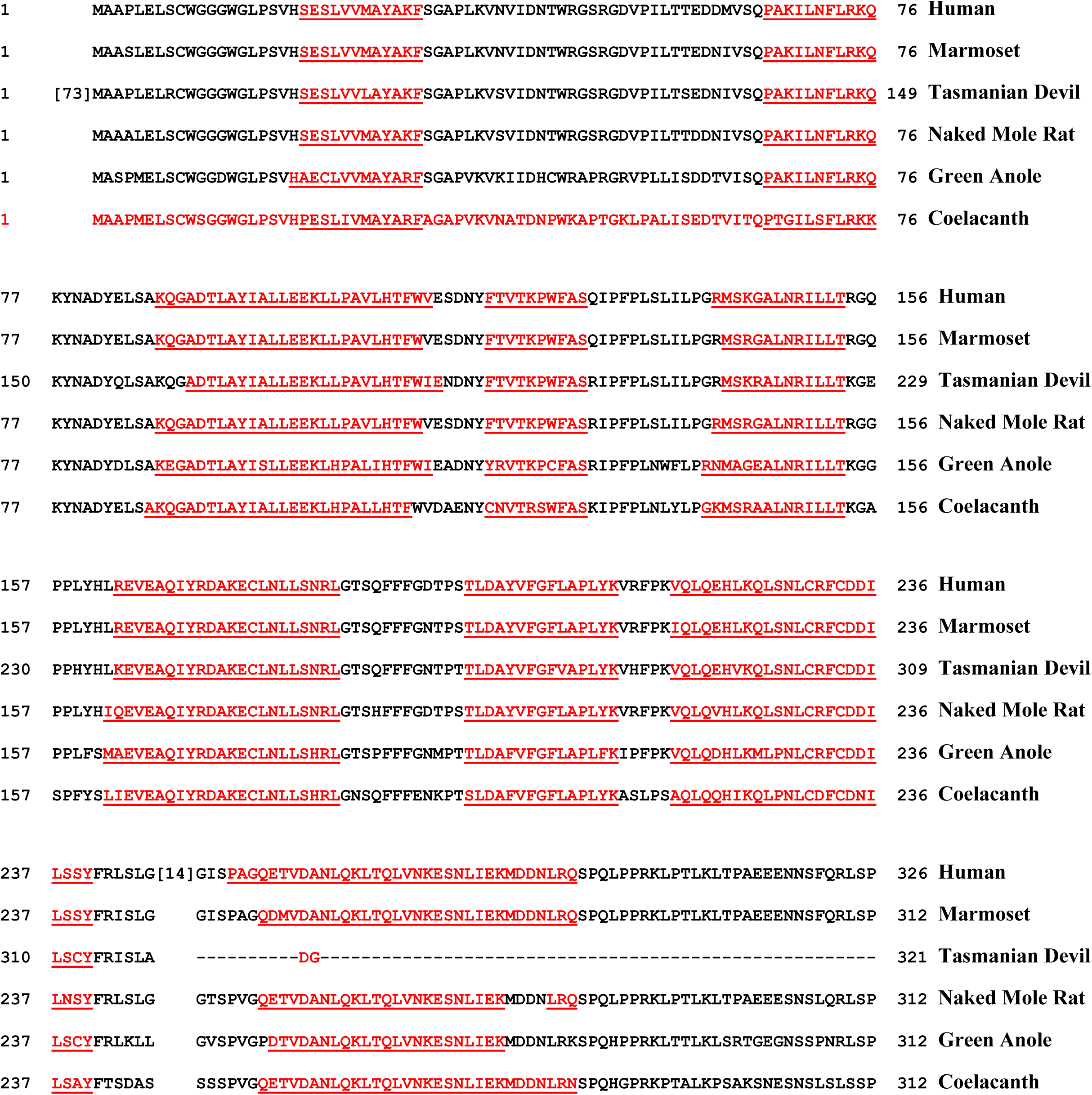
Protein secondary structure of metaxin 3. The figure shows the multiple sequence alignment of selected vertebrate metaxin 3 proteins with predicted alpha-helical segments underlined and in red. The species in the figure represent, from top to bottom: primate, primate, marsupial, rodent, reptile, and fish. Alpha-helical segments are the dominant secondary structure. Little beta-strand was detected. The 9 helical segments are highly conserved among all of the species. The spacings between the segments are also highly conserved. None of the examples, and other species not included, varied from this pattern. Metaxin 1 proteins also possess a similar pattern of 9 helical segments (not shown). Metaxin 2 proteins, with only about 22% amino acid identities compared to metaxin 3, have the same 9 helical segments, plus a 10^th^ helical segment near the N-terminus (not shown). The helical segments were identified using the PSIPRED secondary structure prediction tool (UCL Department of Computer Science).

For alignments such as in Figure 2, the zebrafish metaxin 1a protein sequence was deduced from the metaxin 1a cDNA sequence (GenBank EF202556; K.W. Adolph) determined using Exelixis clone 3545071, derived from a zebrafish testis cDNA library (ZF31XL).

## RESULTS AND DISCUSSION

### Occurrence of Metaxin 3 Proteins

Metaxin 3 genes are highly conserved in the genomes of most mammals, birds, fish, amphibians, and reptiles, as shown by database searches (NCBI). Zebrafish and *Xenopus* metaxin 3 protein sequences, determined from cDNA sequences, were used to identify homologous protein sequences in a wide variety of vertebrates. The conserved, homologous sequences are, generally, amino acid sequences predicted from genomic DNA sequences. In addition to homology, metaxin 3 proteins were identified by the presence of GST_N_Metaxin and GST_C_Metaxin domains, discussed in the next section, but no other major domains other than Tom37. The lengths of the homologous proteins also had to be similar to the lengths of zebrafish and *Xenopus* metaxin 3 proteins (313 and 309 amino acids, respectively).

For invertebrates such as *C. elegans*, sea urchins, and tunicates, metaxin 3 genes have not been detected, although metaxin 1 and 2 genes are present. Plants and bacteria have a single metaxin-like gene that is equally homologous to metaxins 1, 2, and 3 of humans and other higher organisms.

### Domain Structure of Metaxin 3 Proteins

Metaxin 3 proteins are defined by the existence of GST_N_Metaxin and GST_C_Metaxin domains. A Tom37 domain is also a feature of MTX3 proteins. The domain structures of selected MTX3 proteins are shown in Figure 1. For human metaxin 3, the GST_N_Metaxin1_like sequence consists of residues 5 through 77, and the GST_C_Metaxin1_3 sequence of residues 105-241. Tom37 extends between 23 and 142. The upper four domain structures in the figure are those of mammals, including primates (human and bushbaby), a carnivore (Siberian tiger), and a rodent (13-lined ground squirrel). The lower two examples include a reptile (green sea turtle) and a fish (medaka). The high degree of conservation of the metaxin 3 domain structures is apparent in the figure.

Metaxin 1 proteins, such as human MTX1, possess similar domains, in particular GST_N_Metaxin1_like, GST_C_Metaxin1_3, and Tom37 domains. For metaxin 2, the domains are GST_N_Metaxin2 and GST_C_Metaxin2, as well as Tom37.

Glutathione S-transferase proteins also possess N-terminal and C-terminal GST domains, but these are distinct from the GST_Metaxin domains. For example, human GST alpha 1 (222 amino acids) has GST_N_Alpha and GST_C_Alpha domains, human GST mu 1 (218 amino acids) has GST_N_Mu and GST_C_Mu domains, while human GST omega 1 (241 amino acids) has GST_N_Omega and GST_C_Omega domains. Besides having distinctive GST domains, the glutathione S-transferase proteins are smaller than metaxin 3 proteins, such as human metaxin 3 with 326 amino acids.

### Amino Acid Sequence Alignments and Amino Acid Analysis of Metaxin 3 Proteins

The typical results of aligning a metaxin 3 protein sequence with the metaxin 1 and 2 sequences of the same organism are shown in Figure 2. In (A), the amino acid sequence of human metaxin 3 (326 aa) is aligned with that of human metaxin 1 (317 aa). The alignment shows that 45% of the residues are identities and 62% are similarities. The degree of identity is highest in the N-terminal 75% of the proteins, and lowest in the C-terminal 25%. In (B), human metaxin 3 is aligned with human metaxin 2 (263 aa). The overall degree of homology is much lower, with 22% identities and 33% similarities. The low number of identical amino acids is found along the whole length of the proteins. These results demonstrate that human metaxin 3 is a protein distinct from metaxins 1 and 2. In addition, the metaxin 3 gene codes for a protein more homologous to metaxin 1 than 2.

Similar results are found when other metaxin 3 proteins are aligned with metaxin 1 and 2 proteins of the same organism. As examples, mouse metaxins 3 and 1 show 45% identities, while mouse metaxins 3 and 2 show 21%. *Xenopus* metaxins 3 and 1 have 41% identities, compared to metaxins 3 and 2 with 23%. Zebrafish metaxins 3 and 1a have 42% identities, but 3 and 2 have only 22%. There are two metaxin 1 genes in zebrafish, the metaxin 1a gene (GenBank EF202556, submitted by K.W. Adolph) and the metaxin 1b gene (GenBank AY351958, submitted as metaxin 1 by K.W. Adolph). Zebrafish metaxins 1a and 1b share 68% identities.

Fundamental properties of the deduced amino acid sequences of metaxin 3 proteins are given in Table 1. A variety of vertebrates are included. The table compares the numbers of amino acid residues, polypeptide molecular weights, and percentages of acidic, basic, and hydrophobic amino acids. The properties of the sequences are very similar, indicating that the metaxin 3 proteins of very different species are highly conserved.

**Table 1.** Amino Acid Analysis of Metaxin 3 Proteins

Amino Acid Analysis of Metaxin 3 Proteins
| Metaxin 3<br>Source | # Amino<br>Acids | Molecular<br>Weight | % Acidic<br>Residues | % Basic<br>Residues | % Hydrophobic<br>Residues |
| --- | --- | --- | --- | --- | --- |
| Human | 326 | 36,777 | 10.1 | 10.4 | 44.9 |
| Dog | 312 | 35,030 | 10.3 | 10.9 | 45.8 |
| Cat | 312 | 34,998 | 10.3 | 10.9 | 45.6 |
| Elephant | 312 | 34,987 | 10.3 | 10.9 | 45.9 |
| Fruit Bat | 311 | 35,005 | 10.3 | 10.9 | 46.0 |
| Platypus | 292 | 32,744 | 11.0 | 12.0 | 47.2 |
| Guinea Pig | 312 | 34,948 | 10.3 | 9.9 | 45.5 |
| Python | 312 | 35,380 | 10.3 | 11.5 | 44.6 |
| Shark | 308 | 34,915 | 10.1 | 10.4 | 43.4 |

### Genes Adjacent to Metaxin 3 Genes

The genes adjacent to metaxin 3 genes are identical in a wide variety of species, including human, zebrafish, *Xenopus*, dog, shark, elephant, panda, and platypus. As shown in Figure 3A, the order of the genes is: *HOMER1---PAPD4*---*CYMA5*---*MTX3*---*THBS4*---*SERINC5*, where *HOMER1* is homer scaffolding protein 1, *PAPD4* is poly(A) polymerase D4, *CYMA5* is cardiomyopathy associated 5, *MTX3* is metaxin 3 (at 5q14.1 in humans), *THBS4* is thrombospondin 4, and *SERINC5* is serine incorporator 5.

The metaxin 1 gene is also adjacent to a thrombospondin gene, but the other adjacent genes are different: *FAM189B*---*GBA1*---*MTX1*---*THBS3*---*MUC1*---*TRIM46. FAM189B* is family with sequence similarity 189 member B, *GBA1* is glucosylceramidase beta 1 (glucocerebrosidase), *MTX1* is metaxin 1 (at 1q22 in humans), *THBS3* is thrombospondin 3, *MUC1* is mucin1, and *TRIM46* is tripartite motif containing 46. The human genomic region also has *MTX1* and *GBA1* pseudogenes. Figure 3B includes the gene region closest to human metaxin 1. The order is *GBA1*---*psMTX1*---*psGBA1*---*MTX1*---*THBS3*, where *psMTX1* and *psGBA1* are metaxin 1 and *GBA1* pseudogenes.

The metaxin 2 gene region, with *MTX2* at 2q31.1 in humans, is very different, with a group of *Hox* genes adjacent to the *MTX2* gene. A thrombospondin gene is not one of the neighboring genes.

### Phylogenetic Relationships of Metaxin 3 Proteins

The evolutionary relationships of metaxin 3 proteins of a variety of vertebrate species are shown in Figure 4. The phylogenetic tree was derived by multiple sequence alignment of the amino acid sequences. The degree of evolutionary change of the sequences is proportional to the horizontal length of the branches. The results suggest that all of the vertebrate metaxin 3 amino acid sequences are derived from a common ancestor. The analysis also demonstrates that similar vertebrates have similar metaxin 3 amino acid sequences. For example, metaxin 3 sequences of mammals from mice to whales and to primates, including humans, cluster together. The same is true for fish, amphibians, birds, and reptiles. Altogether, 37 vertebrate species showed the pattern of evolutionary relationships illustrated in the figure.

Phylogenetic analysis was also carried out for a number of different vertebrate metaxin 1 and metaxin 2 proteins. The results show that metaxin 3 proteins form a separate cluster distinct from the clusters for metaxin 1 proteins and metaxin 2 proteins. The metaxin 2 cluster shows the greatest divergence, in keeping with the low percentages of identical amino acids in comparing metaxin 2 with 1 and 3. However, the phylogenetic trees indicate that metaxin 3 along with 1 and 2 are likely to share a common ancestral protein sequence.

Invertebrates lack metaxin 3 genes, as mentioned above, although they do possess metaxin 1 and metaxin 2 genes. Plants and bacteria have a single gene equally homologous to metaxins 1, 2, and 3. Because of this, the phylogenetic analysis of metaxin 3 only includes vertebrates.

### Secondary Structures of Metaxin 3 Proteins

The secondary structures of metaxin 3 proteins are dominated by alpha-helices, with little beta-strand. To illustrate this, Figure 5 includes the metaxin 3 proteins of six representative vertebrates. Each is seen to have 9 helical segments with the same pattern of spacing between the segments. The representative metaxin 3 proteins are those of mammals that include primates (human, marmoset), a marsupial (Tasmanian Devil), and a rodent (naked mole rat). Also included in the figure are a reptile (green anole) and a fish (coelacanth). The same pattern of 9 helices is also found for metaxins 1 and 2. Metaxin 2 proteins have, in addition, a 10^th^ helix in their extended N-terminal regions.

A transmembrane helix found near the C-terminus for all metaxin 1 proteins studied is not present in metaxin 3 proteins. For example, human metaxin 1 has a predicted transmembrane helix between residues 272 and 294 of the 317 amino acid protein, but human metaxin 3 does not have a transmembrane helix. A transmembrane helix is also absent in metaxin 2 proteins. The presence of a transmembrane helix is consistent with metaxin 1 proteins being components of the outer mitochondrial membrane. The C-terminal helix could serve to anchor metaxin 1 proteins to the membrane. Metaxin 3 proteins do not therefore seem to be anchored to the outer mitochondrial membrane in the same way.

The possibility of signal peptides at the N-terminus of metaxin 3 proteins was investigated. The results showed that signal peptides are not present. Newly synthesized metaxin 3 proteins are therefore unlikely to be secreted proteins.

